# *Paranannizziopsis* spp. associated with skin lesions in wild snakes in North America and development of a real-time PCR assay for rapid detection of the fungus in clinical samples

**DOI:** 10.1101/2023.07.13.548879

**Authors:** Jeffrey M. Lorch, Megan E. Winzeler, Julia S. Lankton, Stephen Raverty, Heindrich N. Snyman, Helen Schwantje, Caeley Thacker, Susan Knowles, Hugh Y. Cai, Daniel A. Grear

**Affiliations:** U.S. Geological Survey – National Wildlife Health Center, 6006 Schroeder Road, Madison, Wisconsin USA, 53711; Animal Health Centre, Ministry of Agriculture, 1767 Angus Campbell Road, Abbotsford, British Columbia, Canada V3G 2M3; Animal Health Laboratory – Kemptville, University of Guelph, 79 Shearer Street, Kemptville, Ontario, Canada K0G 1J0; Wildlife and Habitat Branch, Ministry of Forests, Lands, Natural Resource Operations and Rural Development, 2080 Labieux Road, Nanaimo, British Columbia, Canada V9T 6J9; Animal Health Laboratory, Laboratory Services Division, University of Guelph, Guelph, Ontario N1G 2W1

## Abstract

The emergence of ophidiomycosis (or snake fungal disease) in snakes has prompted increased awareness of the potential effects of fungal infections on wild reptile populations. Yet, aside from *Ophidiomyces ophidiicola*, little is known about other mycoses affecting wild reptiles. The closely related genus *Paranannizziopsis* has been associated with dermatomycosis in snakes and tuataras in captive collections, and *P. australasiensis* was recently identified as the cause of skin infections in non-native wild panther chameleons (*Furcifer pardalis*) in Florida, USA. Here we describe five cases of *Paranannizziopsis* spp. associated with skin lesions in wild snakes in North America and one additional case from a captive snake from Connecticut, USA. In addition to demonstrating that wild Nearctic snakes can serve as a host for these fungi, we also provide evidence that the genus *Paranannizziopsis* is widespread in wild snakes, with cases being identified in Louisiana (USA), Minnesota (USA), Virginia (USA), and British Columbia (Canada). Phylogenetic analyses conducted on multiple loci of the fungal strains we isolated identified *P. australasiensis* in Louisiana and Virginia; the remaining strains from Minnesota and British Columbia did not cluster with any of the described species of *Paranannizziopsis*, although the strains from British Columbia appear to represent a single lineage. Finally, we designed a pan-*Paranannizziopsis* real-time PCR assay targeting the internal transcribed spacer region 2. This assay successfully detected DNA of all described species of *Paranannizziopsis* and the two potentially novel taxa isolated in this study and did not cross-react with closely related fungi or other fungi commonly found on the skin of snakes. The assay was 100% sensitive and specific when screening clinical (skin tissue or skin swab) samples, although full determination of the assay’s performance will require additional follow up due to the small number of clinical samples (n=14 from 11 snakes) available for testing in our study. Nonetheless, the PCR assay can provide an important tool in further investigating the prevalence, distribution, and host range of *Paranannizziopsis* spp. and facilitate more rapid diagnosis of *Paranannizziopsis* spp. infections that are otherwise difficult to differentiate from other dermatomycoses.

## INTRODUCTION

Mycoses are emerging threats to wildlife conservation and have led to massive population declines and species extinctions (Berger et al. 1998; Skerratt et al. 2007; Cheng et al. 2021). Ectothermic animals, such as amphibians and reptiles, are particularly vulnerable to mycoses because their body temperatures are within a thermal range permissible for the growth of a wide variety of fungi (Fisher et al. 2020). Increased popularity in keeping reptiles as pets beginning in the 1990s (Auliya 2003) facilitated better documentation of fungal infections in lizards, snakes, and turtles. However, aside from chytridiomycosis and ophidiomycosis, fungal infections in wild amphibians and reptiles remain poorly studied.

Of the emerging fungal pathogens affecting reptiles, members of the Onygenales formerly classified as the *Chrysosporium* anamorph of *Nannizziopsis vriesii* (CANV) have received the most attention. Cryptic fungal strains formerly designated as CANV have since been recognized as representing multiple species in at least three genera: *Nannizziopsis, Ophidiomyces*, and *Paranannizziopsis* (Sigler et al. 2013). *Nannizziopsis* includes at least nine species (*N. arthrosporioides, N. barbatae, N. chlamydospora, N. cocodili, N. dermatitidis, N. draconii, N. guarroi, N. pluriseptata*, and *N. vriesii*) known to infect various types of reptilian hosts (Sigler et al. 2013; Stchigel et al. 2013). The sole representative of *Ophidiomyces* (*O. ophidiicola*) is only known to infect snakes (Sigler et al. 2013). Five species of *Paranannizziopsis* are currently recognized (*P. australasiensis, P. californiensis, P. crustacea, P. longispora*, and *P. tardicrescens*); members of this genus have been recovered from snakes, lizards, and tuataras (Sigler et al. 2013; Stchigel et al. 2013; Rainwater et al. 2019; Díaz-Delgado et al. 2020).

The genera *Nannizziopsis, Ophidiomyces*, and *Paranannizziopsis* were originally described from captive animals and are considered emerging pathogens among pet reptiles (Sigler et al. 2013; Mitchell and Walden 2013; Cabañes et al. 2014). *Nannizziopsis guarroi*, the causative agent of yellow fungus disease in captive bearded dragons (*Pogona* spp.) and green iguanas (*Iguana iguana*) (Abarca et al. 2010; Sigler et al. 2013; Gentry et al. 2021), is perhaps the most well-known of the former CANV fungi. The often-fatal disease has been described in captive lizards in North America, Europe, and Asia (Abarca et al. 2010; Sigler et al. 2013; Sun et al. 2021). *Ophidiomyces ophidiicola*, the causative agent of snake fungal disease (SFD) or ophidiomycosis, has similarly been reported from captive snakes in North America, Europe, Asia, and Australia (Sigler et al. 2013; Lorch et al. 2016; Sun et al. 2021). Infections involving *Paranannizziopsis* spp. have been less frequently reported in the literature. In reptiles held in zoological collections, *P. australasiensis* has been isolated from northern tuataras (*Sphenodon punctatus punctatus*) and a bearded dragon (*Pogona barbata*) in New Zealand, file snakes (*Acrochordus* sp.) in Australia, and tentacled snakes (*Erpeton tentaculatum*) and African bush vipers (*Atheris squamigera*) in the USA (Sigler et al. 2013; Díaz-Delgado et al. 2020; Mack et al. 2021). The remaining *Paranannizziopsis* species have each been recovered from single outbreak events primarily involving tentacled snakes (Sigler et al. 2013; Stchigel et al. 2013; Rainwater et al. 2019; Mack et al. 2021).

The natural geographic distributions and effects of the former CANV fungi on wild reptiles is poorly understood. In 2008, *O. ophidiicola* was reported in wild snakes in the USA (Allender et al. 2011). Subsequent studies demonstrated that *O. ophidiicola* had been present in wild snakes in the USA for decades prior to its discovery (Lorch et al. 2021) but was likely introduced from Eurasia (Ladner et al. 2022). *Nannizziopsis barbatae* has been reported in wild lizards in Australia, but it is unclear whether the fungus is native to that continent (Peterson et al. 2020). *Paranannizziopsis australasiensis* was recently reported as a cause of dermatomycosis in an introduced population of wild panther chameleons (*Furcifer pardalis*) in Florida (Claunch et al. 2023). However, we are unaware of any additional verified reports of *Paranannizziopsi*s spp. infections in wild reptiles.

Given that many mycoses of wildlife have emerged through the introduction of exotic fungi into naïve host populations (Drees et al. 2017; O’Hanlon et al. 2018; Ladner et al. 2022), ensuring that reptile-infecting fungi are not translocated and released into areas where they do not naturally occur is important. Thus, better documentation of the distribution and occurrence of these fungi in wild reptiles is critical. Herein, we report cases of *Paranannizziopsis* spp. infections in wild snakes in North America. We confirmed infection using a combination of histopathologic examination, fungus culture, and molecular techniques to detect *Paranannizziopsis* spp. in snakes with skin lesions. We also report a novel real-time PCR assay for *Paranannizziopsis* spp. that could facilitate more rapid and accurate screening for the pathogen.

## METHODS

### Diagnostic Evaluation

Snake samples were submitted from various states in the USA to the U.S. Geological Survey – National Wildlife Health Center (NWHC) or from the lower mainland, Gulf Islands and Vancouver Island, British Columbia to the regional node of the Canadian Wildlife Health Cooperative (CWHC) as part of routine wildlife health surveillance by the Wildlife Health Program, Ministry of Forests, Lands, Natural Resource Operations and Rural Development, British Columbia. All snakes included in this study had gross lesions of skin disease consistent with ophidiomycosis and other dermatomycoses (Lorch et al. 2015).

For histopathologic examination, skin lesions and major internal organs (when an entire carcass was available for examination) were collected, fixed in 10% neutral buffered formalin, processed through a graded series of alcohols to xylene by conventional histologic techniques, embedded in paraffin, sectioned at 5 µm, and stained with hematoxylin and eosin (H&E). In select cases, special stains with periodic-acid Schiff (PAS) and Grocott’s methenamine silver (GMS) were performed to visualize fungal elements. One sample consisted of molted skin only and was not examined histologically. Fresh skin was also collected from each snake for ancillary testing.

Fungus culture was performed as described in Lorch et al. (2016). Briefly, small pieces (c. 2 mm x 2 mm) of fresh lesioned skin were touched several times to different locations on the surface of a dermatophyte test medium (DTM) agar plate, and the skin tissue then placed in the center of the plate. Plates were double-sealed with parafilm and incubated at 30°C. Cultures were checked every two days for a total of 20 days, and fungal colonies representing unique morphotypes were streaked for isolation on fresh DTM plates. Isolated clones were isolated a second time. All fungal isolates were initially identified by sequencing the internal transcribed spacer (ITS) region as described previously (Lorch et al. 2016).

### Multilocus Sequence Analysis

To determine the relationship of strains of *Paranannizziopsis* spp. isolated in this study, we performed a phylogenetic analysis involving multiple loci. In addition to the six novel strains, we included representative strains of all described species of *Paranannizziopsis*: *P. australasiensis, P. californiensis, P. crustacea, P. longispora*, and *P. tardicrescens*; the type strain of *O. ophidiicola* was included as an outgroup (Table S1).

Genomic DNA was obtained from fungal strains grown on Sabouraud dextrose medium containing gentamicin and chloramphenicol at 24°C for a period of 10-21 days. Fungi were ground with glass beads and a pestle to mechanically break the cell wall, and nucleic acid purified using the phenol-chloroform extraction method. In addition to the ITS, portions of seven loci were amplified for use in phylogenetic analyses: actin (*ACT*) gene, β-tubulin (*BTUB*) gene, minichromosome maintenance complex component 7 (*MCM7*) gene, RNA polymerase II second largest subunit (*RPB2*) gene, translation elongation factor 1-α (*TEF*) gene, the D1-D2 domain of the large subunit ribosomal DNA, and the mitochondrial cytochrome oxidase subunit III (*COX3*) gene. PCR was conducted using GoTaq Flexi DNA polymerase (Promega, Madison, Wisconsin, USA) according to the manufacturer’s instructions. Primers and cycling conditions are presented in Table S2. DNA was sequenced in both directions using the same primer pairs used for amplification and internal sequencing primers for large amplicons. All sequences were deposited in GenBank (see Table S1).

Nucleotide sequences were aligned independently for each locus using MUSCLE in MEGA X v10.1.8 (Kumar et al. 2018), and all gaps were deleted. All eight loci were then concatenated for the multigene phylogenetic analyses. A maximum likelihood analysis was performed using IQ-TREE (Minh et al. 2020) through the IQ-TREE web server (Trifinopoulos et al. 2016). Loci were partitioned and the best substitution model for each locus was determined using the “auto” selection feature (Chernomor et al. 2016). The model was run with 1,000 bootstrap alignments and all other settings set to default. For the Bayesian analysis, the best model for each locus was determined using ModelFinder (Kalyaanamoorthy et al. 2017) through the IQ-TREE web server (Trifinopoulos et al. 2016); all other settings were left as default except that candidate models were restricted to those compatible with MrBayes. Best models are shown in Table S1. The Bayesian analysis was performed using MrBayes v3.2.7a (Huelsenbeck and Ronquist 2001; Ronquist and Huelsenbeck 2003) through the CIPRES Science Gateway (Miller et al. 2010). The analysis consisted of two runs with four chains and 5,000,000 generations; the first 25% were discarded as burn-in, and the sampling frequency was set to 1,000 generations.

For some snakes from Canada, *Paranannizziopsis* spp. isolates were not available and tissue samples from which culture could be attempted were not saved. However, extracted DNA from affected skin samples of these animals had been retained. For these samples, tissues were lysed with TriReagent (Thermo Fisher Scientific, Burlington, Ontario, Canada) and nucleic acid extracted using the MagNA 96 instrument with DNA/Viral NA small volume kits (Roche, Laval, Quebec, Canada). *Ophidiomyces ophidiicola* was not detected in these samples by qPCR (Allender et al. 2015) at the Animal Health Laboratory, University of Guelph in Canada.

However, *Paranannizziopsis* spp. were detected in these samples through amplification and sequencing of the ITS region as described above. In addition to the ITS region, *ACT* and *BTUB* loci were amplified directly from these extracts using the same methods described above for isolates. Amplicons were sequenced as described above; those yielding poor quality sequence data (likely due to cross-amplification of DNA from non-target organisms) were cloned using the Zero Blunt TOPO PCR Cloning Kit (Invitrogen, Waltham, Massachusetts, USA) and individual clones sequenced.

### Real-time PCR Assay Development

To develop a pan-*Paranannizziopsis* quantitative PCR, we first aligned ITS sequences for the type strains of *P. australasiensis, P. californiensis, P. crustacea, P. longispora, P. tardicrescens*, and two potentially unique taxa isolated during this study. The presence of numerous indels in the ITS1 made that region unsuitable for primer and probe development. The ITS2 region was more conserved among members of the genus yet divergent from closely related genera (Fig. S1). Primer and probe design was conducted using the PrimerQuest® tool (https://www.idtdna.com/SciTools). The oligo sequences were as follows: forward primer, 5’ – AGCACGGCTTGTGTGTT – 3’; reverse primer, 5’ – CGCAGACACCTGGAACTC – 3’; probe 5’ -(6-FAM) TGGACGGGCCTIAAATGCAGT (BHQ-1) – 3’. An inosine base was placed in the probe sequence due to the presence of a single nucleotide polymorphism in this region among species (Fig. S1). Real-time PCR was conducted using the QuantiNova Probe PCR Kit (Qiagen, Venlo, Netherlands) with each 20-µL reaction consisting of 3.5 µL of water, 10.0 µL of 2x QuantiFast PCR Master Mix, 0.1 µL of QuantiNova ROX Reference dye, 0.4 µL of 20 µM forward primer (0.4 µM final concentration), 0.4 µL of 20 µM reverse primer (0.4 µM final concentration), 0.2 µL of 20-µM probe (0.2 µM final concentration), 0.4 µL of 20-µg/µL bovine serum albumin (0.4 µg/µL final concentration), and 5 µL of DNA template. Cycling was performed with the “fast” setting on an Applied Biosystems® QuantStudio™ 5 Real-time PCR System under the following conditions: 95°C for 2 min; 40 cycles of 95°C for 5 sec, 60°C for 5 sec.

The specificity of the assay was assessed by screening 12 strains of *Paranannizziopsis* spp. (including type isolates of all described species and all strains recovered in this study), 22 strains of closely related onygenalean fungi (“near neighbors,” including *Nannizziopsis* spp. And *O. ophidiicola*, which are known to infect reptiles), and 15 strains of other fungi frequently isolated from the skin of wild snakes (Table S3; Bohuski et al. 2015; Lorch et al. 2016). DNA template from pure cultures of these fungi was extracted using PrepMan™ Ultra Sample Preparation Reagent (Life Technologies, Carlsbad, California, USA) according to the manufacturer’s instructions.

To assess the accuracy of the assay in clinical samples, we extracted DNA from skin tissue or skin swabs of snakes with lesions compatible with dermatomycosis (Table S4). For most of the snakes collected in Canada, DNA was extracted from skin tissue as described above. For one case from Canada (20-3833) and for samples originating in the USA, DNA was extracted using a bead-beating method (Hyatt et al. 2007). For this latter method, samples were extracted using 125 µL of Prepman™ Ultra Sample Preparation Reagent and 100 mg of zirconium beads. Samples were homogenized using an MP Bio FastPrep® (MP Biomedicals, LLC, Irvine, California, USA) on level 6.5 for 45 sec, followed by centrifugation at 13,000 x g for 30 sec. The homogenization steps were repeated, then samples were incubated at 100°C for 10 min, cooled at room temperature for 3 min, centrifuged at 13,000 x g for 3 min, and the supernatant used as PCR template. Clinical samples originated from snakes in which *Paranannizziopsis* spp. were (n=14; some snakes were represented by multiple samples) and were not (n=28) previously detected. Most samples that were *Paranannizziopsis* spp.-negative by culture originated from snakes with ophidiomycosis (Table S4). All samples screened with the *Paranannizziopsis* spp. assay were also screened for the presence of *Ophidiomyces* using a real-time PCR assay targeting the ITS region (Bohuski et al. 2015).

Synthetic double-stranded DNA (gBlocks™ Gene Fragments, Integrated DNA Technologies, Coralville, Iowa, USA) was used as a positive control and to generate standard curves for each run. This positive control was engineered with three nucleotide insertions (see Supplemental) to help distinguish false positive reactions due to contamination of diagnostic samples by the DNA standards. The efficiency of each assay was evaluated by a standard curve, which consisted of a series of seven 10-fold serial dilutions of the synthetic standard ranging from 0.05 fg to 50,000 fg of synthetic DNA per reaction (or approximately 1.1 × 10^2^ – 1.1 × 10^8^ copies per reaction) using the QuantStudio™ Design and Analysis software (version 1.5.1, Life Technologies Corporation). Each standard was performed in triplicate on each plate. For each PCR run that included diagnostic samples from snakes, a negative extraction control (extraction buffer without a swab added) and a nuclease-free water control were included.

To determine the limits of detection (LOD) and quantification (LOQ) for the assay, 5 µL of synthetic DNA template ranging from 7.62 × 10^−6^ fg to 10,000 fg (approximately 0.172 – 224,874,408 copies) per reaction were each run 80 times on 10 separate plates (12 dilution series per plate; Table S5). The threshold was set to 4% of the maximum fluorescence, and any sample that crossed the threshold on or before 45 cycles was considered positive. We determined the LOD using these replicate dilution series and a hierarchical logistic regression where the detection of target synthetic DNA was determined as a Bernoulli distributed variable (π) with probability as a linear function of synthetic DNA concentration in reaction *i*, run on plate *j. Y*_*i,j* was the observed detection in reaction *i* from plate *j*, and *Xi*_ was the concentration of synthetic DNA in reaction *i*. We included parameters to account for the possibility of false-negative (*S*) or false-positive (*C*) results independent of the target concentration. We specified these as closed-form informative priors but did not estimate them under our experimental design. The linear functional form of probability of detection was specified with random effects intercept and slope estimated for each separate plate (α_*j*_ and β_*j*_, respectively) that was estimated as normally distributed parameters from an overall intercept (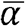) and slope (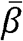). This allowed for propagation of uncertainty for inter-plate variation into the estimation of detection as a function of concentration,

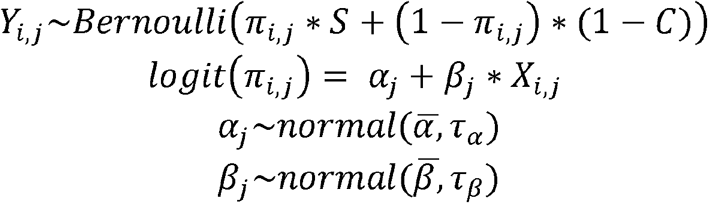

With closed form informative priors for base false-negative and false-positive rates,

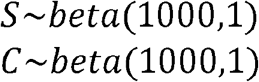

And non-informative hyper-priors for regression coefficients

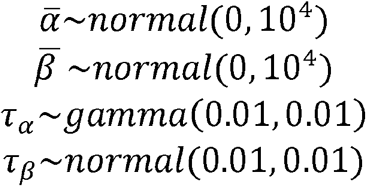

We made inference using MCMC and Gibbs sampling implemented in rJAGS using 50,000 interactions with 20% burn-in using three independent MCMC chains with randomly selected starting values from each parameter’s prior distribution (Plummer 2016). We assessed MCMC convergence visually and by using the Gelman-Rubin statistic with the five MCMC chains to ensure that the scale reduction factor for each parameter was <1.1 (Brooks and Gelman 1998). After discarding the burn-in, we combined the three MCMC chains and sampled every 10^th^ iteration for the posterior estimation of the parameters. We then used the posterior draws of 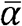 and 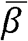 to calculate the mean and 95% Highest Posterior Density Interval (95% HPD) LOD as concentration of synthetic DNA detection with 95% probability as,

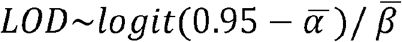

To determine the quantification curve and LOQ, we used a hierarchical log-linear regression following Sivaganesan et al. (2008) that incorporates plate to plate variability, similar to the detection model. The cycle threshold (Ct) of detection of target synthetic DNA was determined as a normally distributed mean (*µ*) with a linear relationship to the log_10_ synthetic DNA concentration (*X*) in reaction *i*, run on plate *j*. The linear functional form of probability of detection was specified with random effects intercept and slope estimated for each separate plate (*α*_*j*_ and *β*_*j*_, respectively) that were estimated as normally distributed parameters from an overall intercept (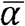) and slope (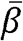).

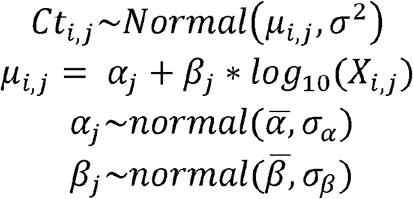

And non-informative hyper-priors for regression coefficients

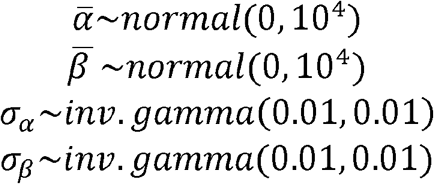

We implemented parameter estimation using MCMC and Gibbs sampling and the same procedures as the detection parameter fitting. The LOQ was derived by calculating the relative error of the estimated mean copy number per reaction, *n*, for each standard dilution series quantity, *k*.

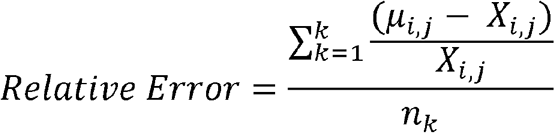

The LOQ was set as the lowest quantity where the relative error of all estimates to that quantity was <25%.

## RESULTS

### Diagnostic Evaluation

The study cohort consisted of eight whole snake carcasses submitted for necropsy that had died or were euthanized prior to submission, two skin punch biopsies collected from live snakes, and a molted skin from a live snake (Table 1). One sample (the molted skin; case number NWHC 46748-1) originated from a snake in a captive collection in which multiple animals were described as having skin lesions. The remaining five samples from the USA represented wild snakes for which single animals from a given location were noted as having clinical signs of skin infection. Samples from Canada represented a possible cluster of cases from southern British Columbia involving gartersnake species (*Thamnophis* spp.). Of the eight carcasses examined, one snake (NWHC 26922-5) was part of a mortality event of unknown cause and had a very mild infection that was not thought to be related to the cause of death (Terrell et al. 2020), another snake had evidence of trauma that was likely the cause of death (NWHC 26609-1), and the six snakes from British Columbia were euthanized due to extensive crusting skin lesions.

**Table 1:** List of snakes examined for this study.

| Case Number | Host | Location | Sample Type | Host Status | Collection Date | *Paranannizziopsis* sp. | Detection Method |
| --- | --- | --- | --- | --- | --- | --- | --- |
| 20-3833 | *Thamnophis* sp. | Gabriola Island, British Columbia, Canada | carcass | wild | 10 June 2020 | *Paranannizziopsis* sp. 2 | culture, qPCR |
| NWHC 26609-1 | *Pituophis catenifer* | Chisago County, Minnesota, USA | carcass | wild | 16 June 2015 | *Paranannizziopsis* sp. 1 | culture, qPCR |
| NWHC 27236-1 | *Pantherophis* sp. | Fairfax County, Virginia, USA | biopsy | wild | 22 April 2016 | *P. australasiensis* | culture, qPCR |
| NWHC 24878-7 | *Coluber constrictor* | Virginia Beach County, Virginia, USA | biopsy | wild | 26 April 2014 | *P. australasiensis* | culture, qPCR |
| NWHC 26922-5 | *Liodytes rigida rigida* | Orleans County, Louisiana, USA | carcass | wild | 5 November 2015 | *P. australasiensis* | culture, qPCR |
| NWHC 46748-1 | *Gonyosoma boulengeri* | Connecticut, USA | molted skin | captive | 18 March 2018 | *P. australasiensis* | culture, qPCR |
| K22-047521A | *Thamnophis elegans* | Blue Heron Reserve, Chilliwack, British Columbia, Canada | carcass | wild | 25 May 2022 | *Paranannizziopsis* sp. 2 | panfungal PCR, qPCR |
| K22-047521B | *Thamnophis elegans* | Blue Heron Reserve, Chilliwack, British Columbia, Canada | carcass | wild | 25 May 2022 | *Paranannizziopsis* sp. 2 | panfungal PCR, qPCR |
| K22-047523 | *Thamnophis elegans* | Blue Heron Reserve, Chilliwack, British Columbia, Canada | carcass | wild | 25 May 2022 | *Paranannizziopsis* sp. 2 | panfungal PCR, qPCR |
| K22-053463 | *Thamnophis sirtalis* | Hayward Lake, British Columbia, Canada | carcass | wild | 21 June 2022 | *Paranannizziopsis* sp. 2 | panfungal PCR, qPCR |
| K22-056652 | *Thamnophis elegans* | Lighthouse Reserve, Vancouver Island, British Columbia, Canada | carcass | wild | 30 June 2022 | *Paranannizziopsis* sp. 2 | panfungal PCR, qPCR |

Snake carcasses from the USA exhibited skin lesions consisting primarily of few to many raised white to brown areas of discoloration or crusting on the head and body (Fig. 1A). In the subset of samples from Canada, skin lesions ranged in size from 1-3 mm up to 3-6 mm in diameter among the individual snakes. The lesions were well circumscribed, raised, thickened, occasionally roughened or nodular, and randomly distributed throughout the dorsal, dorsolateral, lateral and, to a much lesser extent, midventral aspect of the body (Fig. 1B). In more severely affected areas, the scales were raised, discolored dark brown to black, and the trailing margins were roughened. These lesions were nonspecific and similar to those reportedly caused by other etiologies, including *O. ophidiicola* (Jacobson 1980; Lorch et al. 2015). Based on the presence of body fat at necropsy, six snakes were assessed as being in fair to good body condition and two snakes were considered to be in poor body condition.

**Fig 1:**
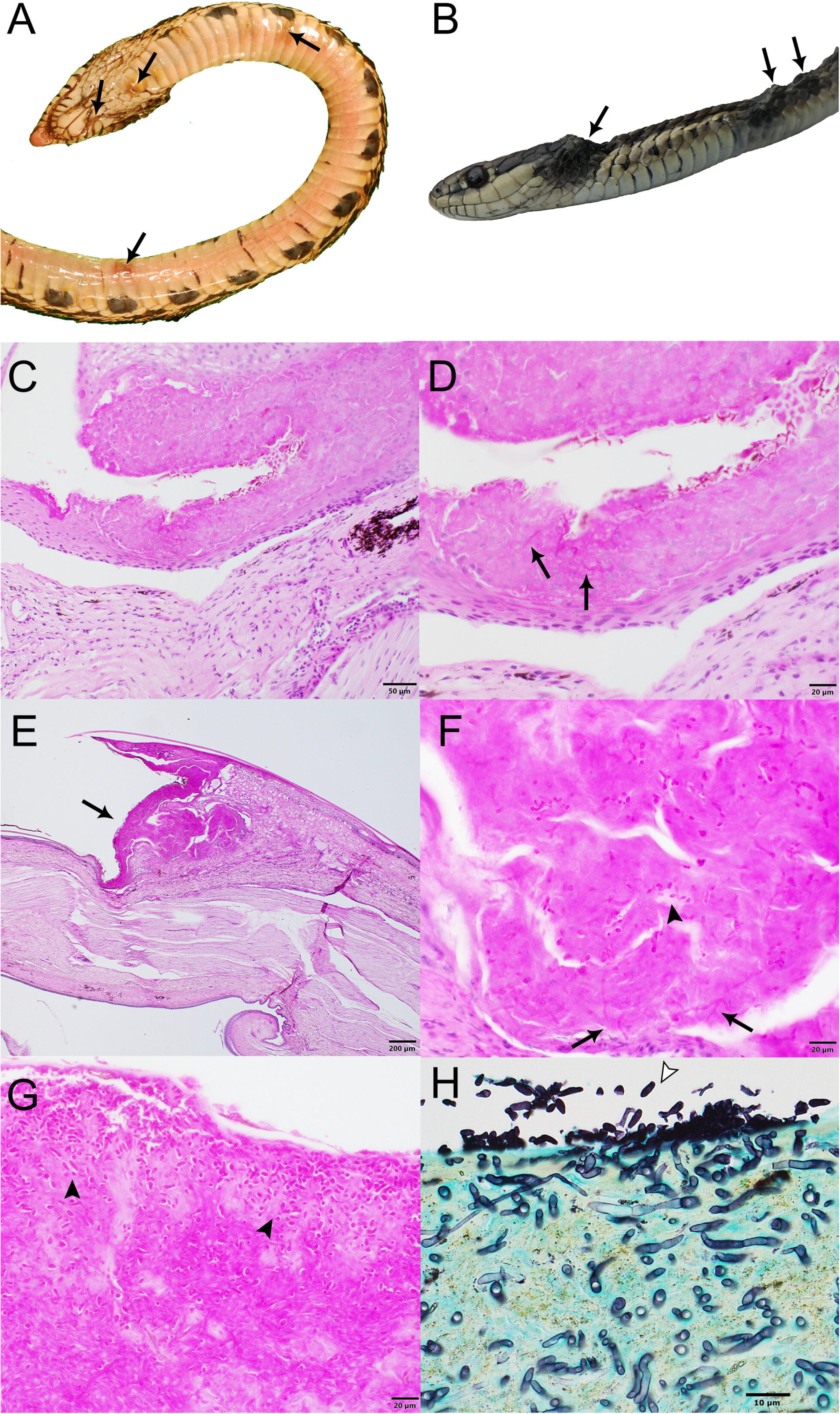
Gross and histopathological images of select wild snakes with *Paranannizziopsis* spp. infections. (A) Ventral surface of gophersnake (*Pituophis catenifer*) infected with *Paranannizziopsis* sp. 1 (NWHC 26609-1). Arrows indicate representative gross lesions. (B) gartersnake (*Thamnophis* sp.) infected with *Paranannizziopsis* sp. 2 (20-3833). Arrows indicate representative gross lesions. (C,D) Glossy crayfish snake (*Liodytes rigida rigida*) infected with *P. australasiensis* (NWHC 26922-5) (PAS stain; scale bar, 50 µm). (C) Areas of epithelial necrosis extend to the deep portions of the epidermis (PAS stain; scale bar, 20 µm). (D) Detail of C showing an area of necrotic epithelium containing low numbers of ∼2-µm diameter faintly PAS-positive fungal hyphae (arrows) (PAS stain; scale bar, 200 µm). (E-G) Same gophersnake as in (A) (PAS stain). (E) Within the crypt between two scales is a large area of epithelial necrosis (arrow) (scale bar, 200 µm). (F) Moderate numbers of ∼2-x 5-µm ovoid aleurioconidia (arrowhead) and ∼ 2-µm diameter fungal hyphae (arrows) are within the area of necrosis (scale bar, 20 µm). (G) Within a crust overlying the epithelium are many ∼2-x 5-µm ovoid aleurioconidia (arrowheads) (scale bar, 20 µm). (H) Western terrestrial gartersnake (*Thamnophis elegans*) infected with *Paranannizziopsis* sp. 2 (K22 2022-047521A). Extensive full thickness necrosis and ulceration was present along the jaws and head of this snake with the necrotic debris being widely permeated by a meshwork of silver positive fungal hyphae. An area of epithelial necrosis contains many superficial ovoid to bullet-shaped aleurioconidia (arrowhead) and deep fungal hyphae is shown (GMS stain; scale bar, 10 µm).

Six samples examined yielded *Paranannizziopsis* spp. in culture, and *Paranannizziopsis* ssp. was detected in the remaining five samples by panfungal PCR of the ITS region. Internal transcribed spacer region DNA sequences of the isolates revealed that strains for four snakes most closely matched the type strain of *P. australasiensis* (99.6% identity). A fifth strain (hereafter referred to as *Paranannizziopsis* sp. 1) isolated from a wild snake in Minnesota had an ITS sequence that most closely matched *P. tardicrescens* (98.3% identity), while a sixth strain (hereafter referred to as *Paranannizziopsis* sp. 2) from wild snakes in British Columbia had an ITS sequence that most closely matched *P. californiensis* (97.5% identity).

Snakes for which skin was collected from carcasses or by biopsy punches were examined by light microscopy; the molted skin sample was not examined. Histopathologically, all snakes exhibited epidermal necrosis with intralesional fungal hyphae ranging from 2-5 µm in diameter (Fig. 1). Two of the three snakes examined histopathologically from which *P. australasiensis* was isolated (NWHC 24878-7; NWHC 26922-5) had necrotizing granulocytic epidermitis and heterophilic to mononuclear dermatitis (Fig. 1C, 1D). A biopsy from the third snake (NWHC 27236-1) with *P. australasiensis* consisted only of epidermis, and crush artifacts in the epidermis made interpretation difficult, although mild epidermitis was suspected. In addition to fungal hyphae, ovoid or bullet-shaped ∼2 × 5 µm aleurioconidia were within necrotic epidermis in two of these three snakes (NWHC 24878-7; NWHC 27236-1). Microscopic lesions observed in the snakes with *P. australasiensis* indicated a primary fungal etiology and were consistent with previous descriptions of *Paranannizziopsis* spp. infections (Bertelsen et al. 2005; Rainwater et al. 2019). One snake (NWHC 24878-7) had both arthroconidia (∼2 × 4 µm in size) and ovoid aleurioconidia present in the skin section.

The snake from which *Paranannizziopsis* sp. 1 (NWHC 26609-1) was cultured had epidermal necrosis, erosion, and (rarely) ulceration with a myriad of intralesional fungal hyphae and bacteria (Fig. 1E-1G). Fungal hyphae were highly variable in size (1-6 µm in diameter), indicating that multiple fungal species may have been present. Ovoid to bullet-shaped ∼2-x 5-µm aleuroconidia were often present within areas of necrotic epidermis. Inflammation associated with fungal hyphae in the epidermis or dermis was minimal.

The snakes from which *Paranannizziopsis* sp. 2 was detected (20-3833; K22-047521A, K22-047521B; K22-047523; K22-053463; K22-056652) had variably extensive full-thickness necrosis and ulceration of the epidermis and scales with replacement by abundant thick serofibrinous exudate, eosinophilic necrotic debris, and variable infiltrates of degenerate and necrotic heterophils, macrophages, and fewer lymphocytes. Diffusely permeating the serocellular debris and extending to inflammatory bands were a myriad of clear to faintly basophilic hyaline fungal hyphae. The hyphae were thin-walled, pale, parallel, regularly to irregularly septate, and had a dichotomously right angle and occasionally haphazardously branching pattern. Clusters of 3-to 4-µm ovoid aleurioconidia were present superficially (Fig. 1H; Table S5). Unique to infections associated with *Paranannizziopsis* sp. 2 was fungal invasion into the dermis and inflammation in underlying skeletal musculature in association with myocellular degeneration and necrosis. In one snake, necrosis extended into the mandibular bone.

### Multilocus Sequence Analysis

The final alignment consisted of 5,278 characters (*ACT*: 753; *BTUB*: 866; *COX3*: 611; D1-D2: 571; *ITS*: 447; *MCM7*: 571; *RPB2*: 828; *TEF*: 631). Trees resulting from the maximum likelihood and Bayesian analyses both shared identical topologies. Similar to previous phylogenetic studies, we found that the genus *Paranannizziopsis* consists of two core clades, with the first clade (Clade I) consisting of *P. australasiensis, P. californiensis*, and *P. tardicrescens*, and the second clade (Clade II) containing *P. crustacea* and *P. longispora* (Fig. 2). Consistent with the ITS sequence results, four of the newly recovered strains (NWHC 24878-7, NWHC 26922-5, NWHC 27236-1, and NWHC 46748-1) formed a well-supported subclade with two described strains of *P. australasiensis* (UAMH 10439, UAMH 11645). There was no sequence variation among the strains in this subclade within the portions of the D1-D2, *MCM7, RPB2*, or *TEF* loci that were examined. However, at least some of the newly isolated strains from snakes in North America differed from UAMH 10439 and UAMH 11645 by 1-2 SNPs in sequences of the ITS, *ACT, BTUB*, and *COX3* loci. Strains UAMH 10439 and UAMH 11645 were 100% identical to one another at all loci examined, indicating clonality. However, the newly isolated strains from North American snakes, while often having identical haplotypes at a given locus, each had unique genotypes when all loci were analyzed together. Two additional novel strains (*Paranannizziopsis* sp. 1 and *Paranannizziopsis* sp. 2) resided within Clade I but were divergent from described species of *Paranannizziopsis* (Fig. 2). No strains isolated in this study were members of Clade II.

**Fig 2:**
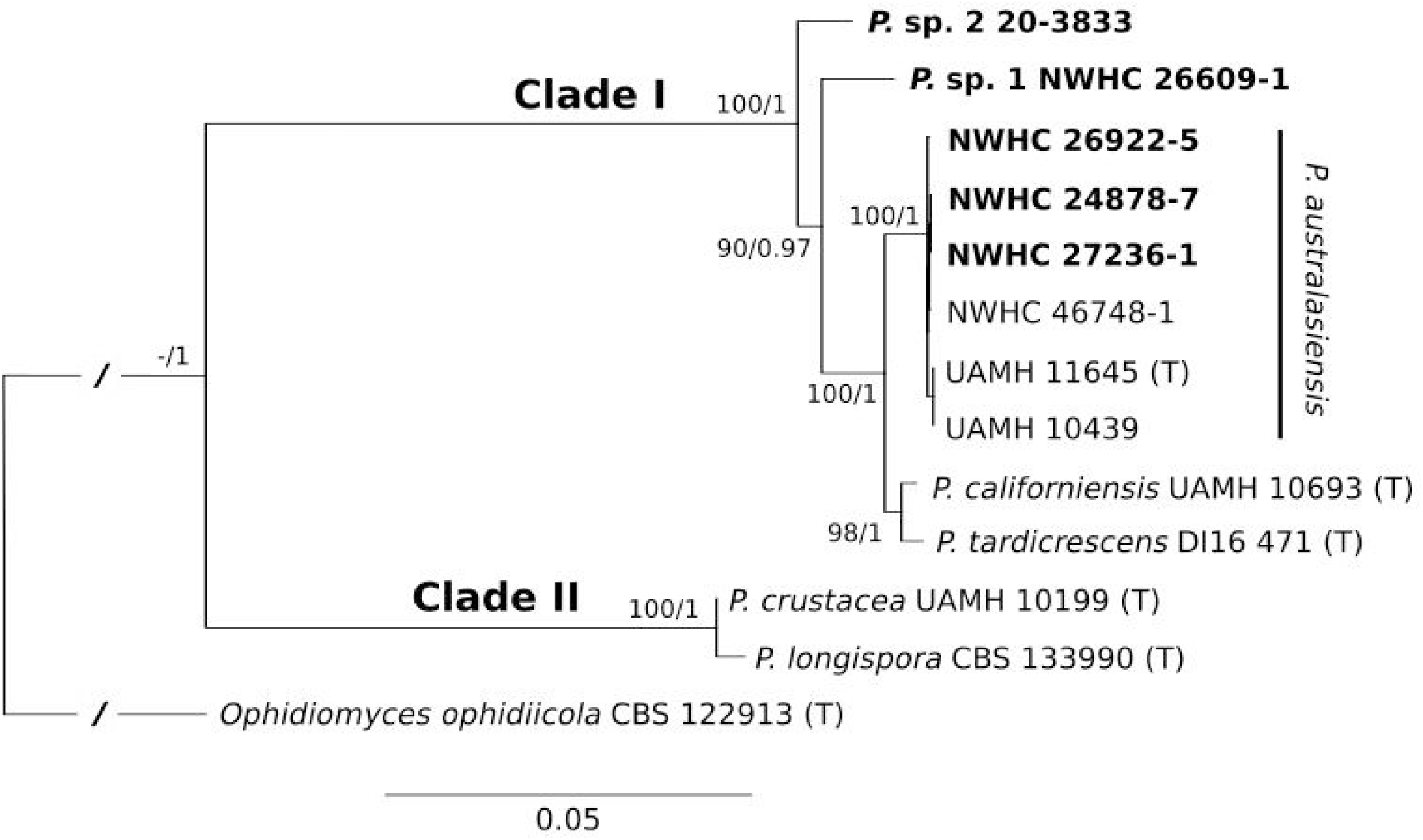
Best tree resulting from a maximum likelihood (ML) analysis of eight concatenated loci of *Paranannizziopsis* spp. strains. The consensus tree from a Bayesian analysis had an identical topology. Boldened strains are those isolated from wild snakes in this study. Two of these strains resided within the *P. australasiensis* clade but were not identical to existing strains belonging to that species. The remaining two strains did not cluster with described taxa and may be novel species. Support values are presented at each node (bootstrap support values from the ML analysis/posterior probabilities from the Bayesian analysis). Support values within the *P. australasiensis* clade are not shown due to space. Type strains are designated with “(T)”. The tree is rooted with *Ophidiomyces ophidiicola*. Branch lengths for the root have been shortened (indicated by “/”) to make the tree easier to view.

In addition to the *Paranannizziopsis* spp. isolates, we successfully sequenced the ITS from affected skin of five snakes from British Columbia for which no isolates were available. We also were able to sequence *ACT* and *BTUB* loci from four and three of these animals, respectively. We targeted these loci because they exhibited both inter and intraspecific variation; thus, they can provide information on whether strains of the *Paranannizziopsis* infecting snakes in British Columbia likely represent a single species. Sequences generated from these four snakes were all 100% identical to those of *Paranannizziopsis* sp. 2.

### Real-time PCR Assay Development

The diagnostic assay performed in the commonly accepted range of standard curve efficiency between 90% and 110% (Life Technologies 2011). The mean efficiency was 96.682% with an average coefficient of determination (R^2^ of 0.999). The LOD was 7.74 copies per reaction (1.548 copies per µL) with a 95% confidence interval of (5.63, 11.12) copies per reaction (Fig. S2; Table S5). The LOQ of the assay was determined to be 13.7 copies per reaction (2.75 copies per µL) and corresponded to a Ct value of ∼37.

Based on screening of DNA extracted from pure cultures, the sensitivity and specificity of the *Paranannizziopsis* spp. real-time PCR assay were both 100%. That is, all isolates belonging to described species of *Paranannizziopsis*, as well as *Paranannizziopsis* sp. 1 and *Paranannizziopsis* sp. 2, showed evidence of amplification for the assay whereas other fungi within Onygenales or commonly associated with snake skin did not generate detectable PCR product. When the assay was applied to clinical samples, the sensitivity and specificity were both 100%. Two samples (∼22%; cases NWHC 24878-7 and NWHC 26922-5) that were PCR-positive for *Paranannizziopsis* spp. were also PCR-positive for *O. ophidiicola*. Case 20-3833 was weakly positive (Ct = 38.17) for *Ophidiomyces* by real-time PCR when tested at an outside laboratory; however, two samples from this snake tested at the NWHC were PCR-negative for *O. ophidiicola* (Table S4).

## DISCUSSION

*Paranannizziopsis* spp. have been repeatedly associated with morbidity and mortality in captive reptiles on multiple continents (Sigler et al. 2013; Stchigel et al. 2013; Rainwater et al. 2019; Díaz-Delgado et al. 2020; Mack et al. 2021) as well as in a population of wild non-native panther chameleons in the USA (Claunch et al. 2023). The cases described herein represent the first confirmed instances (to our knowledge) of *Paranannizziopsis* spp. infections in wild snakes and expand the known distribution of the pathogen in wild reptiles in the USA. Specifically, we isolated *P. australasiensis* from three wild snakes in Louisiana (n=1) and Virginia (n=2), USA. Two additional isolates from wild snakes in Minnesota (USA) and British Columbia (Canada) were genetically divergent from described species of *Paranannizziopsis*.

The prevalence of *Paranannizziopsis* spp. infections in wild reptiles in North America and its effects on populations would benefit from further study. Detections from distant locations on the continent would seemingly indicate that the fungus is geographically widespread.

However, cases of *Paranannizziopsis* spp. infection in North America are relatively infrequent given the large number of wild snakes that have been assessed and diagnosed with dermatitis due to ophidiomycosis (Lorch et al. 2016). This could indicate that *Paranannizziopsis* spp. rarely acts as a consequential pathogen in Nearctic snake species. Indeed, we were unable to attribute *Paranannizziopsis* spp. as a primary cause of significant disease in some of the snakes we examined, especially those infected with *P. australasiensis*. Likewise, several outbreaks in captive snakes were concomitant with other health issues (Bertelsen et al. 2005; Rainwater et al. 2019; Díaz-Delgado et al. 2020; Mack et al. 2021), making it difficult to determine directionality of *Paranannizziopsis* spp. infection as either a cause or consequence of these other disease processes. Nonetheless, the cases of infection with *Paranannizziopsis* sp. 2 that we documented in British Columbia were more severe and appeared to be primary in nature. This could indicate that prevalence and disease outcomes vary geographically, by host species, or by the species of *Paranannizziopsis* that an animal is infected with. The paucity of *Paranannizziopsis* spp. infections in wild snakes in North America could be due to the previous lack of diagnostic tools to aid in pathogen detection (e.g., *Paranannizziopsis* spp.-specific PCR), biases in pathogen screening (e.g., screening primarily for *O. ophidiicola* in snakes with skin lesions), or highly localized occurrences of the fungus (perhaps due to recent introduction).

The origin of *Paranannizziopsis* species found on wild snakes in North America is unclear. Two isolates of *P. australasiensis* recovered from separate outbreaks at zoological parks in Australia and New Zealand were 100% identical to one another at all eight loci we examined. An additional three isolates from captive animals in New Zealand were also 100% identical to these based on available ITS sequences in GenBank (sequence data for the other seven loci were not available). These findings indicate that a single clonal lineage was likely involved in the Australasian outbreaks. In contrast, isolates of *P. australasiensis* from the USA were more genetically diverse, and none were identical to one another at all loci examined. These findings could be compatible with a pathogen having been established in an area for a long period of time. The infrequent detection of *Paranannizziopsis* spp. in Nearctic reptiles could be the result of inapparent clinical signs of infection (e.g., host tolerance or resistance) due to co-evolution of the host and pathogen; under this hypothesis, the manifestation of disease in reptiles originating from Africa, Asia, and Australasia could be the result of the pathogen coming into contact with naïve host species.

An alternative explanation is that species of *Paranannizziopsis* represent introduced fungal pathogens in North America. Given the frequency with which reptiles are imported for the pet trade, multiple introductions of *Paranannizziopsis* spp. to North America, as has been described for *O. ophidiicola* (Ladner et al. 2022), cannot be ruled out. In captive reptiles, *Paranannizziopsis* spp. have predominantly been recovered from host species native to southeast Asia, Australia, or New Zealand. This is notable because many of these host species (e.g., file snake, tentacled snake, rhinoceros snake [*Gonyosoma boulengeri*], Wagler’s viper [*Tropidolaemus wagleri*]) are, or until recently were, primarily obtained as wild-caught (rather than captive-bred) individuals that may have been harboring *Paranannizziopsis* spp. at the time of collection. Given that most cases of *Paranannizziopsis* spp. infection in captive reptiles have occurred in institutions that likely harbor many reptile species, it is difficult to determine which hosts are responsible for introducing the fungus into a collection. Efforts to sample wild reptiles around the world could help determine where the fungus originates.

We do not attempt to describe the potentially novel *Paranannizziopsis* species recovered during this study because each was isolated from a single snake (although *Paranannizziopsis* sp. 2 was detected by molecular means on multiple snakes). With the exception of *P. australasiensis*, material used to describe all other species of *Paranannizziopsis* originated from single outbreak events within an institution or collection. Strains recovered during such outbreaks are likely clonal in nature. As mentioned above, *P. australasiensis* may also have been described from clonal strains despite being isolated from different zoological collections. Species descriptions based on recent clonal lineages do not capture the genetic diversity or morphological variation that exists within a taxon. Our multilocus phylogenetic analysis indicates that some named species of *Paranannizziopsis* exhibit minimal genetic divergence (e.g., *P. crustacea* and *P. longispora*; *P. californiensis* and *P. tardicrescens*; Fig. 2). Additional sampling and characterization of strains from regions where these fungi occur naturally, analyses to look for recombination between taxa, and mating experiments would be useful to better define species boundaries and determine whether there has been taxonomic oversplitting within the genus.

The majority of cases of fungal dermatitis in snakes in the USA is caused by *O. ophiodiicola* (Lorch et al. 2016). For research and diagnostic purposes, SFD status is often inferred on the basis of clinical signs or clinical signs in conjunction with detection of *O. ophiodiicola* (Hileman et al. 2018; Baker et al. 2019). However, caution is warranted in assigning SFD status, especially when diagnostic accuracy is important. Real-time PCR assays are often used to confirm the presence of *O. ophiodiicola* on snakes with skin lesions (Allender et al. 2015; Bohuski et al. 2015). Yet, these assays are so sensitive that they frequently detect low levels of the fungus on the skin of snakes without clinical signs (e.g., Hileman et al. 2018; McKenzie et al. 2019). Four of the 10 cases of *Paranannizziopsis* spp. infection in wild snakes that we investigated occurred in areas where SFD is endemic (Lorch et al. 2016). Thus, *O. ophiodiicola* could conceivably occur on the skin of snakes that have lesions caused by *Paranannizziopsis* spp., leading to a false diagnosis of SFD. Indeed, we simultaneously detected *O. ophidiicola* and *P. australasiensis* in skin lesions of two of four (50%) wild snakes that originated from an SFD endemic area in this study. In one of these cases (NWHC 26922-5), we tested two lesions from the same snake; *O. ophidiicola* DNA was detected at low levels in only one of the lesions while *Paranannizziopsis* spp. was detected in both lesions. In the second snake (NWHC 24878-7), *O. ophidiicola* DNA was also detected at low levels in the single lesion that was sampled. Furthermore, only *P. australasiensis* was recovered in culture from these two snakes, indicating that *O. ophidiicola* may have been unrelated to the disease process. A low level of *O. ophidiicola* DNA was detected on a snake from British Columbia by an outside laboratory, but we were unable to replicate this finding when testing two separate skin lesions from the same animal.

Distinguishing between ophidiomycosis and *Paranannizziopsis* spp. infection can also be challenging based on histopathology. We did not observe any pathognomonic microscopic lesions that would help differentiate the two infections. The presence of arthroconidia is often considered to be characteristic of *O. ophiodiicola* infections (Baker et al. 2019). However, we do not always observe arthroconidia in ophidiomycosis cases. Furthermore, arthroconidia were reported in histologic sections of snakes infected with *P. australasiensis* and *P. crustacea* (Mack et al. 2021; Claunch et al. 2023), and *P. tardicrescens* produces arthroconidia in culture (Rainwater et al. 2019). In most cases of *Paranannizziopsis* spp. infection we examined, arthroconidia were not observed in histopathology. The one exception was a snake (NWHC 24878-7) from which *O. ophidiicola* DNA was detected in low quantities, and we cannot rule out that this snake was co-infected with *O. ophidiicola* and *P. australasiensis*. The presence or absence of arthroconidia should not be considered a reliable characteristic for distinguishing ophidiomycosis from *Paranannizziopsis* spp. infections, and the potential for co-infections complicates diagnoses. Revision of the existing case definition for ophidiomycosis (Baker et al. 2019) is warranted in light of these findings. However, a larger sample set of cases of *Paranannizziopsis* spp. infection need to be reviewed to better document the characteristics of the infection and how it differs from SFD. The interactions between *O. ophidiicola* and *P. australasiensis* are not known, and experimental co-infection trials could help disentangle whether the pathogens compete with one another or facilitate more severe infections on their hosts.

The novel qPCR assay described herein may help overcome some diagnostic challenges by providing a tool to rapidly detect nucleic acid of *Paranannizziopsis* spp. in clinical samples. Specifically, this assay exhibited a 100% diagnostic sensitivity and specificity rate when screening fungal isolates or clinical samples and had an estimated LOD of 18.83 copies per µL and LOQ of 42.16 copies per µL of the target sequence. The LOQ allows us to quantify the amount of target sequence at low copy numbers and provide presence/absence data below the LOQ. Given that the ITS region exists as multiple copies on the fungal genome, the assay can likely detect and quantify very few cells of the target organism within a sample. We intentionally designed the *Paranannizziopsis* spp. qPCR to be broad and detect all currently known species within the genus; potentially novel strains that are close relatives of the described species are also likely to be detected by the assay. Although the assay does not directly speciate the type of *Paranannizziopsis* detected in a sample, follow up sequencing of amplicons can assist with identification. Specifically, *Paranannizziopsis* sp. 1 and sp. 2 possess unique SNPs within the amplified region, and Clade II (*P. crustacea* and *P. longispora*) can be separated from the sublineage of Clade I consisting of *P. australasiensis, P. californiensis*, and *P. tardicrescens* based on one SNP (see Fig. S1). Due to the limited amount of clinical material available from reptiles with *Paranannizziopsis* spp. infections, the qPCR assay could be improved by additional validation, especially for assessing false negative rates and applications beyond clinical samples (e.g., screening environmental samples). However, the assay may prove useful in further investigations into the geographic and host range of *Paranannizziopsis* spp., as well as aiding in clinical diagnoses.

Reptiles as a group would benefit from conservation (Cox et al. 2022). Yet relatively little is known about infectious diseases affecting reptiles and their broader effects on populations. Specifically, many infectious diseases of reptiles have been reported only in captivity; the occurrence and severity of these pathogens both within and outside their areas of natural distribution have not been documented. Although our detections of *Paranannizziopsis* spp. in wild snakes in North America represent an important contribution to the field of reptile disease, a better understanding of the ecology and significance of this pathogen to native populations of reptiles in North America would be beneficial.

## DATA AVAILABILITY

Data associated with this study are available at https://doi.org/10.5066/P9TIGBUU (Lorch et al., 2023).

## ACKNOWLEDGEMENTS

We thank Dr. Adrian Walton (Dewdney Animal Hospital) and the members of the public for reporting and collecting affected snakes in British Columbia and numerous field personnel and state agencies for submitting samples for analysis in the USA. We also thank Dr. Valerie Shearn-Bochsler (U.S. Geological Survey – National Wildlife Health Center) for contributing a case included in this manuscript. We are indebted to staff at the U.S. Geological Survey – National Wildlife Health Center, Animal Health Centre, and Animal Health Laboratory for assistance with processing samples, histopathology, and conducting microbiology and molecular biology analyses. We thank Dr. Nathan Wiederhold (UT Health San Antonio) for generously providing the type strain of *P. tardicrescens*. Any use of trade, firm, or product names is for descriptive purposes only and does not imply endorsement by the U.S. Government.

