## Supplemental Text for "*Paranannizziopsis* spp. associated with skin lesions in wild snakes in North America and development of a real-time PCR assay for rapid detection of the fungus in clinical samples"

This file includes:

Supplemental Text

Fig. S1

Fig. S2

Fig. S2

Table S1 legend

Table S2 legend

Table S3 legend

Table S4 legend

Table S5 legend

DISCLAIMER: Any use of trade, firm, or product names is for descriptive purposes only and does not imply endorsement by the U.S. Government.

SUPPLEMENTAL TEXT

A synthetic double-stranded gBlock™ gene fragment was used as a positive control for real-time PCR and to generate the standard curves for the real-time PCR assay described in this study. The fragment sequence is based on that of the ITS2 region of the type strain of *Paranannizziopsis australasiensis* (UAMH 11645), but has three base pair insertions (shown shaded in gray) added outside the primer and probe binding sites (underlined) to help distinguish the synthetic fragment from DNA of *Paranannizziopsis* spp. Note that the sequence extends beyond both primer binding locations to account for possible DNA degradation at the 3’ and 5’ ends of the gene fragment.

GCACATTGCGCCCCCTGGCATTCCGGGGGGCATGCCTGTCCGAGCGTCATTGCACCCCTCAAGCACGGCTTGTGTGTTGGGCCGCTGTCCCCCGTGGACGGGCCTCAAATGCAGTGGCGTTGCGCCCGAGTTCCAGGTGTCTGCGCGCATGGGAAGCCATCGCCGCGAGACCCGGTCGGTGCCCGTCCGGTTGAACCATTGTCCTT

SUPPLEMENTAL FIGURES

Fig. S1: Alignment of the portion of internal transcribed spacer region 2 (ITS2) of *Paranannizziopsis* species targeted by the *Paranannizziopsis* spp. real-time PCR assay. Single nucleotide polymorphisms relative to the *P. australasiensis* reference sequence are bolded and indicated by an asterisk (*). Note that only a single representative of *P. australasiensis* is presented as all strains of *P. australasiensis* examined shared 100% sequence identity in this region. The primer and probe sequences are presented above the alignment; arrows indicate directionality from the 5’ to 3’ ends of the oligos.

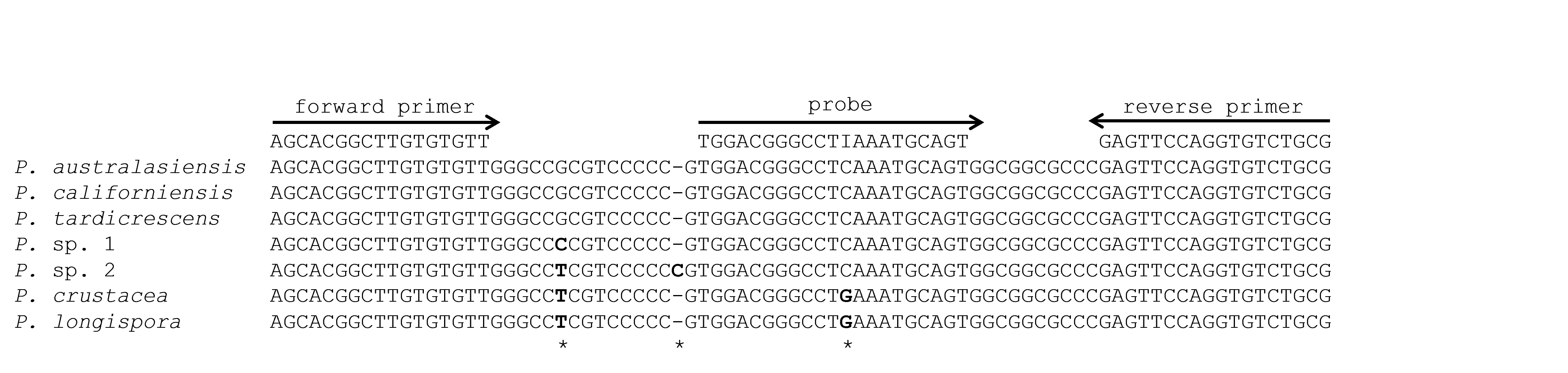

Fig S2: The limit of detection of the *Paranannizziopsis* spp. real-time PCR assay was determined by running multiple replicates of serial dilution of synthetic target DNA and estimating target copy number where detection occurred with 95% confidence (dotted line).

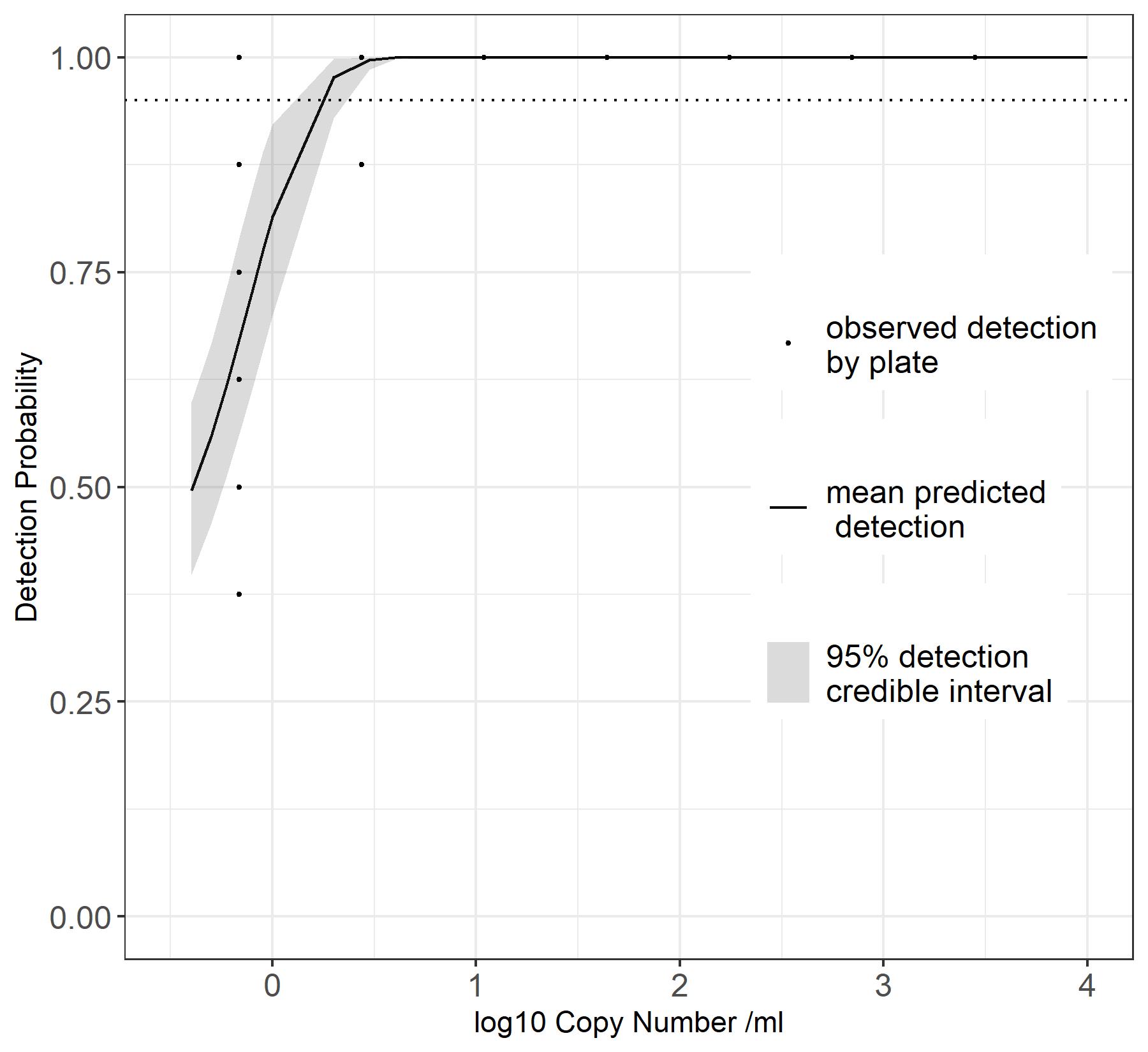

Fig S3: Quantification curve (a) and limit of quantification (b, inset) of the *Paranannizziopsis* spp. real-time PCR assay was determined by running multiple replicates of serial dilutions of synthetic target DNA; the limit of quantification was determined as the lowest dilution DNA template dilution (copy number) where the relative error of all estimates of that dilution quantity was <25%.

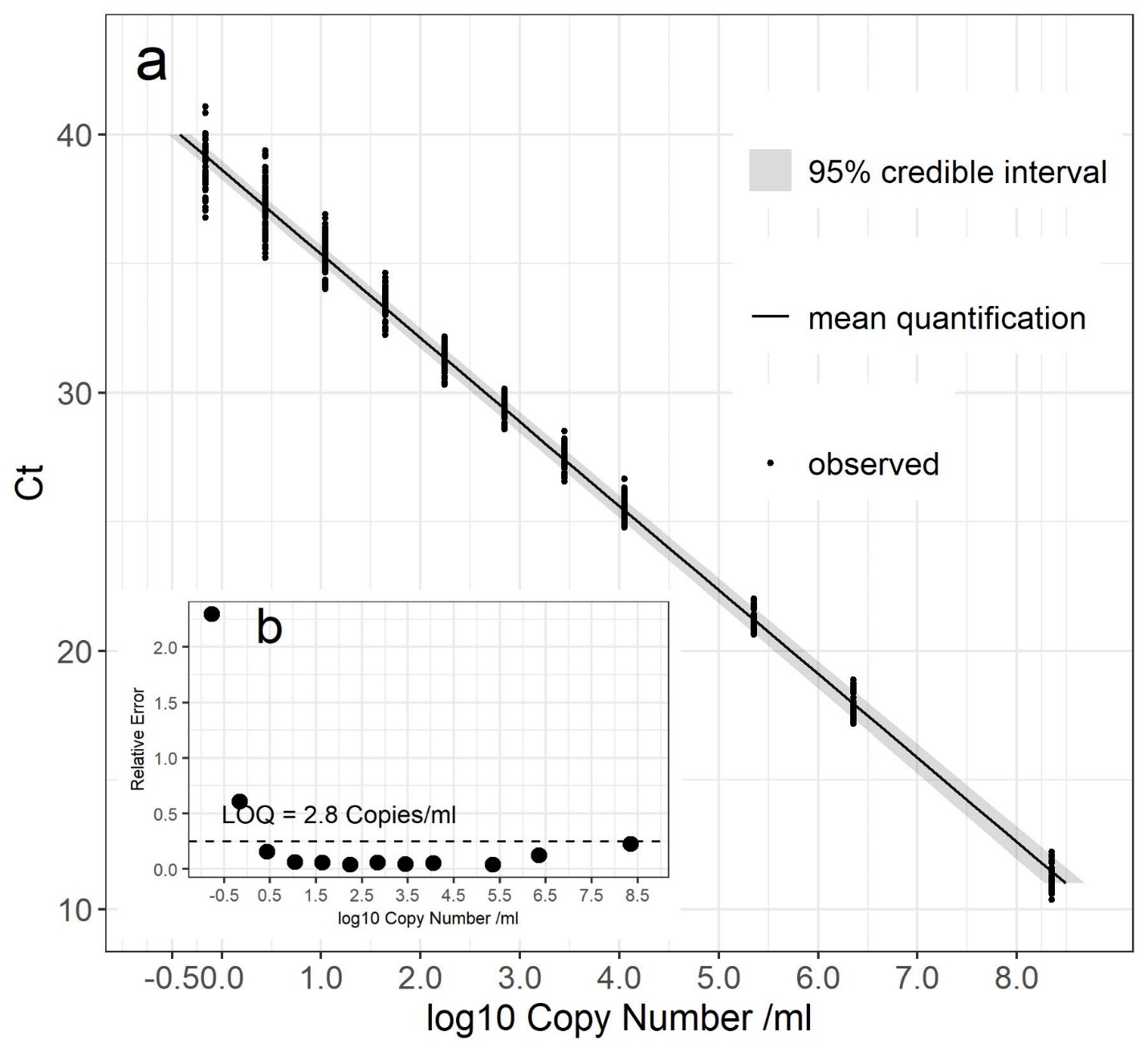

SUPPLEMENTAL TABLE LEGENDS

Table S1: GenBank numbers for loci used for the multigene phylogenetic analyses (provided as a separate file).

Table S2: List of primers and cycling conditions used for amplification of loci for the multigene phylogenetic analyses (provided as a separate file).

Table S3: Fungal isolates used for validation of the *Paranannizziopsis* spp.-specific real-time PCR assay. Strains were categorizes as “target organism” (i.e., *Paranannizziopsis* sp. that should generate a positive PCR result), “near neighbor non-target” (i.e., onygenalean fungus closely related to the genus *Paranannizziopsis* that should not yield a positive PCR result), or “snake-associated non-target” (i.e., fungus that is frequently detected on snake skin, but is not closely related to the genus *Paranannizziopsis*). (provided as a separate file).

Table S4: Snake skin swab samples from which extracted DNA was used for validation of the *Paranannizziopsis* spp.-specific real-time PCR assay. All samples originated from snakes with skin lesions. The samples were screened for the presence of both *Ophidiomyces ophidiicola* and *Paranannizziopsis* spp. by specific real-time PCR. “*Paranannizziopsis* status” was determined based on the previous detection of *Paranannizziopsis* sp. by fungus culture or panfungal PCR from which the snake the swab sample was derived. (provided as a separate file).

Table S5: Serial dilution and real-time PCR detection data of synthetic DNA template used to calculate limit of detection and limit of quantification of the *Paranannizziopsis* spp.-specific real-time PCR assay. The serial dilutions were run on 12 plates. The quantity of the target template (synthetic double-stranded DNA) is reported in fg; target copy numbers per reaction and the resulting cycle threshold value (Ct) are listed. (provided as a separate file).
